# Targeted biallelic integration of an inducible Caspase 9 suicide gene for safer cellular therapies prevents development of drug-resistant escapees in human iPSCs

**DOI:** 10.1101/2021.09.19.460940

**Authors:** Stephanie Wunderlich, Alexandra Haase, Sylvia Merkert, Kirsten Jahn, Maximillian Deest, Silke Glage, Wilhelm Korte, Andreas Martens, Andreas Kirschning, Andre Zeug, Evgeni Ponimaskin, Gudrun Goehring, Mania Ackermann, Nico Lachmann, Thomas Moritz, Robert Zweigerdt, Ulrich Martin

**Affiliations:** Leibniz Research Laboratories for Biotechnology and Artificial Organs (LEBAO), Department of Cardiothoracic, Transplantation and Vascular Surgery, Hannover Medical School, Hannover, Germany; REBIRTH - Research Center for Translational Regenerative Medicine; Department of Psychiatry, Social Psychiatry, and Psychotherapy, Hannover Medical School, Hannover, Germany; Institute for Laboratory Animal Science, Hannover Medical School, Hannover, Germany; Institute for Organic Chemistry, Leibniz University Hannover, Hannover, Germany; Department of Cellular Neurophysiology; Hannover Medical School, Hannover, Germany; Department of Human Genetics, Hannover Medical School, Hannover, Germany; Department of Pediatric Pneumology, Allergology and Neonatology, Hannover Medical School, Hannover, Germany; RG Reprogramming and Gene Therapy, Hannover Medical School, Hannover, Germany; Institute of Experimental Hematology, Hannover Medical School, Hannover, Germany; JRG Translational Hematology of Congenital Diseases, Hannover Medical School, Hannover, Germany

## Abstract

Drug-inducible suicide systems may help to minimize risks of cellular therapies due to the tumor forming potential of human induced pluripotent stem cells (hiPSCs). Recent research challenged the usefulness of such systems since rare drug-resistant subclones were observed that showed elimination or silencing of the transgene.

We have introduced a drug-inducible Caspase9 suicide system (iCASP9) into the AAVS1 safe harbor locus of hiPSCs. In these cells, apoptosis could be efficiently induced *in vitro*. In mice, drug treatment generally led to rapid elimination of teratomas, but individual animals subsequently formed tumor tissue from monoallelic iCASP9 hiPSCs. Very rare drug-resistant subclones of monoallelic iCASP9 hiPSCs appeared *in vitro* with frequencies of ~ 3×10^-8^. Transgene elimination, presumably via Loss of Heterozygosity (LoH), or methylation of the CAG promoter but not methylation of the ppp1r12c locus were identified as underlying mechanisms. In contrast, we never observed any escapees from biallelic iCASP9 hiPSCs, even after treatment of up to 0.8 billion hiPSCs.

In conclusion, biallelic integration of an iCASP9 system in the AAVS1 locus may substantially contribute to the safety level of iPSC-based therapies, which should be calculated by relating clonal escapee frequencies to the cell number in tumors of a size that is readily detectable during routine screening procedures.

## Introduction

Pluripotent stem cell (PSC) technologies come out of age and a number of clinical trials applying embryonic stem cell (ESC) or induced pluripotent stem cell (iPSC)-based cell products are ongoing or in preparation. The tumorigenic potential of PSC-derived cell products, however, either in terms of contaminating undifferentiated cells that form teratoma or due to malignant transformation caused by mutations acquired and enriched during reprogramming ^1^ or culture expansion ^2^ is considered as a major safety concern.

While improved protocols for targeted differentiation of PSCs into the therapeutic derivatives of interest substantially reduced the risk for teratoma formation, introduction of synthetic fail-safe systems with drug-inducible suicide genes have been proposed to further decrease tumour risks.

Such fail-safe systems include the inducible expression of the herpes simplex virus-thymidine kinase (HSV-TK), which has been clinically applied already more than 25 years ago ^3^, and the inducible Caspase9 (iCASP9) safety switch system ^4^ that has been clinically applied more recently in T cells using a retroviral approach ^5^.

In a first study that attempted to quantitatively define the safety level of PSC transplants with integrated fail-safe systems, ESCs carrying an HSV-TK system controlled by one allele of the endogenous CDK1 locus were applied ^6^. Despite general functionality of the HSV-TK suicide system concerning elimination of cycling ESCs, rare proliferating subclones of mouse ESCs were observed *in vivo* and *in vitro* with an average frequency of 6.6×10^-8^ that became resistant to ganciclovir. Remarkably, not only transgene silencing but also loss of heterozygosity (LoH) was identified as underlying mechanism for failure of the integrated drug-inducible suicide system in mouse ESCs. Finally, the authors demonstrated that in case of ESCs with biallelic (homozygous) integration of HSV-TK in the CDK1 locus, no resistant clones appeared among a total of 1.2×10^8^ cells pointing to a substantially higher safety level.

One potential disadvantage of the applied HSV-TK system is that HSV-TK as viral transgene may be immunologically recognized *in vivo* leading to killing of HSV-TK expressing cells even without application of ganciclovir. It should be stressed that placing a HSV-TK transgene under control of a cell cycle-dependent locus such as CDK1 (cyclin-dependent kinase 1) ^6^, indeed restricts the targeted elimination to cycling cells such as tumour cells. This concept, though, is probably not applicable if the iPSC-derived graft has to proliferate in order to fulfil its therapeutic function, because all cycling cells that express the HSV-TK gene may become target of the host’s immune response. In addition, the HSV-TK fail-safe system is especially effective in fast-dividing cells and may fail to eliminate populations of slowly dividing tumour cells. Also, the use of the prodrug ganciclovir as potent antiviral drug e.g. for treatment of serious herpes virus infections may not be possible in affected patients.

Because of these limitations and the lack of data for human iPSCs (hiPSCs), we aimed to quantitatively define the safety level of hiPSC transplants with another integrated drug-inducible suicide system, the iCASP9 safety switch. In this system, the Caspase activation and recruitment domain of human Caspase 9, a gene expressed in fetal and adult tissue that is upregulated during induction of apoptosis, has been replaced by a modified dimerizer-binding domain of the FK506-binding protein 12 ^5^. This system requires caspase dimerization after application of an otherwise bioinert small-molecule (Chemical Inducer of Dimerization, CID) for activation of intracytoplasmic caspase-3 /7 and induction of apoptosis ^4,7^ (**Figure 1A**). Because Caspase 9 is an endogenous gene, it is considered non-immunogenic.

**Figure 1:**
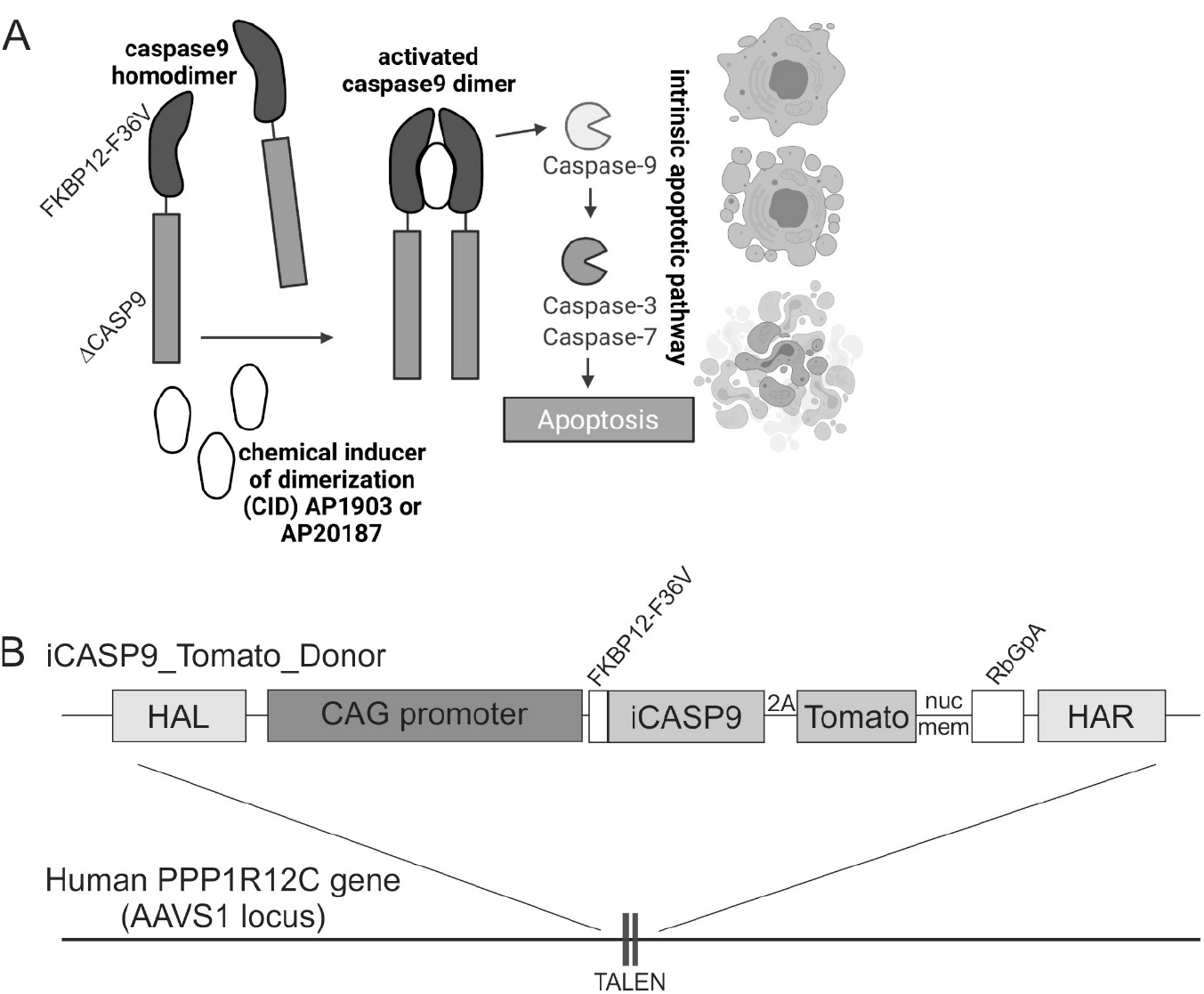
The iCASP9 suicide switch and TALEN based integration into the safe harbor locus AAVS1 in human iPSCs. A) Function of the iCASP9 suicide system. The CASP9 gene is deleted for its endogenous caspase activation and recruitment domain (ΔCASP9) and was coupled to the sequence of a mutated FK506-binding protein, FKBP12, with an F36V mutation that confers increased affinity for chemical inducers of dimerization (CID). FKBP12-F36V binds to otherwise bioinert small-molecule dimerizing agents AP1903 and AP20187, which are lipid-permeant tacrolimus analogues. In the presence of the drugs, the iCASP9 pro-molecule dimerizes and activates the intrinsic apoptotic pathway, leading to cell death via apoptosis. (Scheme adapted from Gargett and Brown, Pharmacology 2014 p.1 Article 235). B) Schematic illustration of iCASP9 donor construct and the AAVS1 target site. The iCASP9 donor construct consists of HAL and HAR left and right handed homology arms, a CAG promoter, an iCASP9 gene coupled via a 2A site to a dTomato fluorescence marker with a nuclear membrane localization signal. Integration of the iCASP9 donor construct was achieved via TALEN based gene editing into the AAVS1 located in the PPP1R12C gene on human chromosome 19.

In order to ensure safe application of the safety switch and efficient cell killing also in slowly dividing hiPSC derivatives we have integrated the iCASP9 construct into the the PPP1R12C / AAVS1 (Adeno-Associated-Virus Site 1) safe harbour locus ^8^. iCASP9 was placed under control of a synthetic promoter consisting of the cytomegalovirus early enhancer element and chicken beta-actin (CAG) promoter, known for robust expression in undiffentiated hiPSCs and their differentiated progeny ^9,10^ (**Figure 1B**). In consideration of the results of Liang et al. ^6^ we have generated hiPSC clones either heterozygous or homozygous for the integrated iCASP9 safety switch. Potential emergence of cell clones escaping the induced suicide was monitored *in vivo* and *in vitro* among therapeutically relevant cell numbers.

## Results

### Generation of Human iPSC Lines with heterozygous or homozygous (mono- or biallelic) integration of an iCASP9 suicide gene coupled to a Tomato fluorescent reporter

In order to enable inducible induction of apoptosis, we applied transcription activator-like effector nucleases (TALEN)-mediated gene editing to integrate an iCASP9 safety switch into the AAVS1 safe harbor locus. iCASP9 gene was place under control of the CAG promoter. A dTomato-fluorescent protein fused to a nuclear membrane localization signal (dTomato_nucmem_) was utilized to facilitate indirect monitoring of iCASP9 expression and discrimination from autofluorescence, and was coupled to the iCASP9 gene via a 2A site (**Figure 1B**). Allele-specific PCR analysis confirmed site-specific integration into one or both alleles of the AAVS1 locus of Phoenix ^11^ and iCBPSC2 ^12^ lines (**Figure S1A&B**). For all further procedures and analyses, one correctly targeted monoallelic (heterozygous) and biallelic (homozygous) clone per cell line was chosen. These clones showed considerable levels of dTomato_nucmem_ expression (**Figure 1C**), absence of chromosomal aberrations as demonstrated by karyotyping (**Figure 1D**) expressed typical pluripotency markers, differentiated into derivatives of all three germ layers and formed teratoma upon injection into NOD*SCID* mice (**Figure S2**). For clarity, all selected iCASP9 cell lines that were applied thereafter are abbreviated according to their origin and genotype: monoallelic iCASP9 Phoenix, biallelic iCASP9 Phoenix, monoallelic iCASP9 hCBiPS2, biallelic iCASP9 hCBiPS2.

### Chemical inducers of dimerization (CIDs) efficiently induce apoptosis in monoallelic and biallelic iCASP9 iPSC clones

Undifferentiated cells of monoallelic and biallelic iCASP9 Phoenix and iCASP9 hCBiPS2 clones cultivated as monolayer on geltrex were treated for 24h with different concentration of two CID variants AP1903 and AP20187 before staining of viable cells with Calcein staining. Intercalating DNA-binding live-dead dyes such as 7-AAD or propidium iodide that were used in previous studies ^10,13^ were found not to be useful for this purpose apparently because dead cells after DNA degradation are detected by flow cytometry as false negative “viable” cells. We were never been able to confirm such seemingly viable cells after reseeding in cell culture (data not shown). In contrast, Calcein provided much more reliable results. The the non-fluorescent acetomethoxy derivate of calcein is transported through the cellular membrane into live cells. Intracellular esterases that are lacking in dead cells remove the acetomethoxy group, the molecule gets trapped inside and gives out strong green fluorescence. Determination of the proportion of Calcein^pos^ cells by FACS revealed very efficient killing of cells of all four mono-/bi-allelic cell lines with both CID variants at concentration ≥ 0,1nM (**Figure 2**).

**Figure 2:**
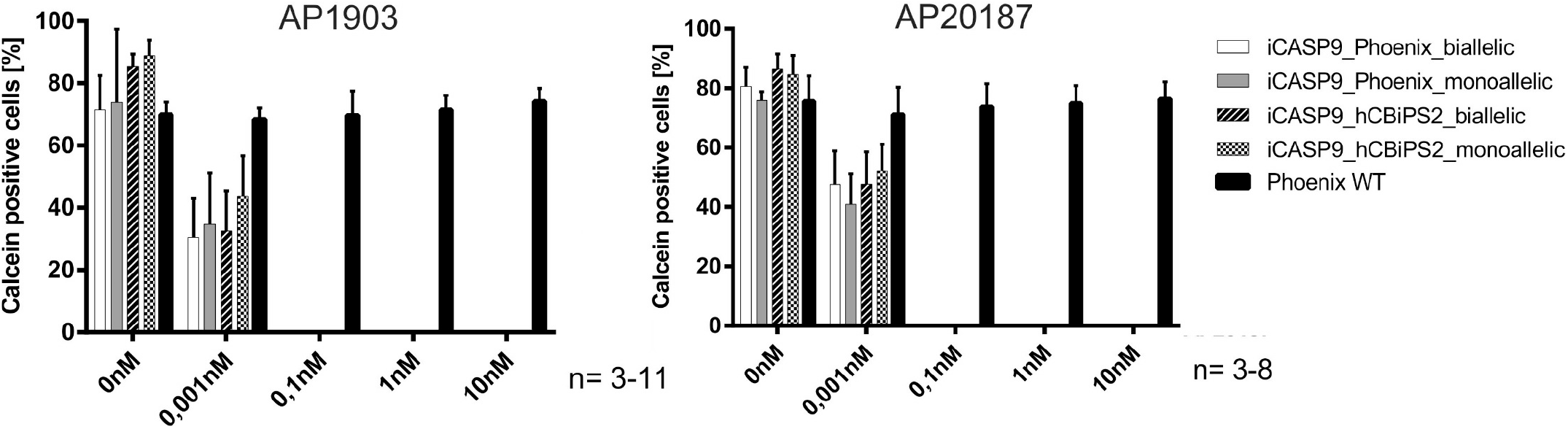
Chemical inducers of dimerization (CIDs) efficiently induce apoptosis in monoallelic and biallelic iCASP9 iPSC clones. FACS analysis of undifferentiated iCASP9 clones cultivated on geltrex with or without CID AP1903 / AP20187 treatment for 24h. As demonstrated by loss of Calcein viability staining of iCASP9 iPSCs, efficient cell killing is achieved already by application of 0.1 nM AP1903 or AP20187 (mean ± SEM; n=3-11).

### *In vivo* CID treatment of preformed teratoma after injection of monoallelic iCASP9 hiPSCs eliminates human cells and teratoma in most but not all mice

*In vivo* experiments in NODSCID mice were conducted to confirm the above results *in vivo* (**Figure 3A**). All mice after injections of iPSCs under the kidney capsule showed massive increase of girth as typical sign for teratoma formation after 8 weeks. Mice that received non transgenic Phoenix wildtype cells and subsequent CID treatment, as well as mice after injection of monoallelic iCASP9 Phoenix iPSCs followed by vehicle-treatment all developed teratoma that stained positive for human nuclear antigen (**Figure 3B**). In contrast, in three out of five mice injection of transgenic monoallelic iCASP9 Phoenix iPSCs followed by treatment with CID resulted in almost complete elimination of frequently cystic teratoma structures (**Table 1** and **Figure 3B**). In these mice, only some abnormal puffy mouse tissue with eosinophilic infiltration around the kidney could be detected (upper middle panel), probably representing fibrotic mouse tissue that formed in response to the massive CID-induced cell death and the resulting infiltration of phagocytes and granulocytes followed by pro-fibrotic cytokine release. In the remaining two out of five mice also extensive elimination of frequently cystic teratoma structures was observed. However, in these mice also tumor like tissue was detected around the kidney that stained positive with an anti-human nucleoli antibody, suggesting that despite killing of the vast majority of engrafted cells, re-growths of escapees occurred that were apparently resistant to CID-induction of iCASP9-mediated suicide.

**Figure 3:**
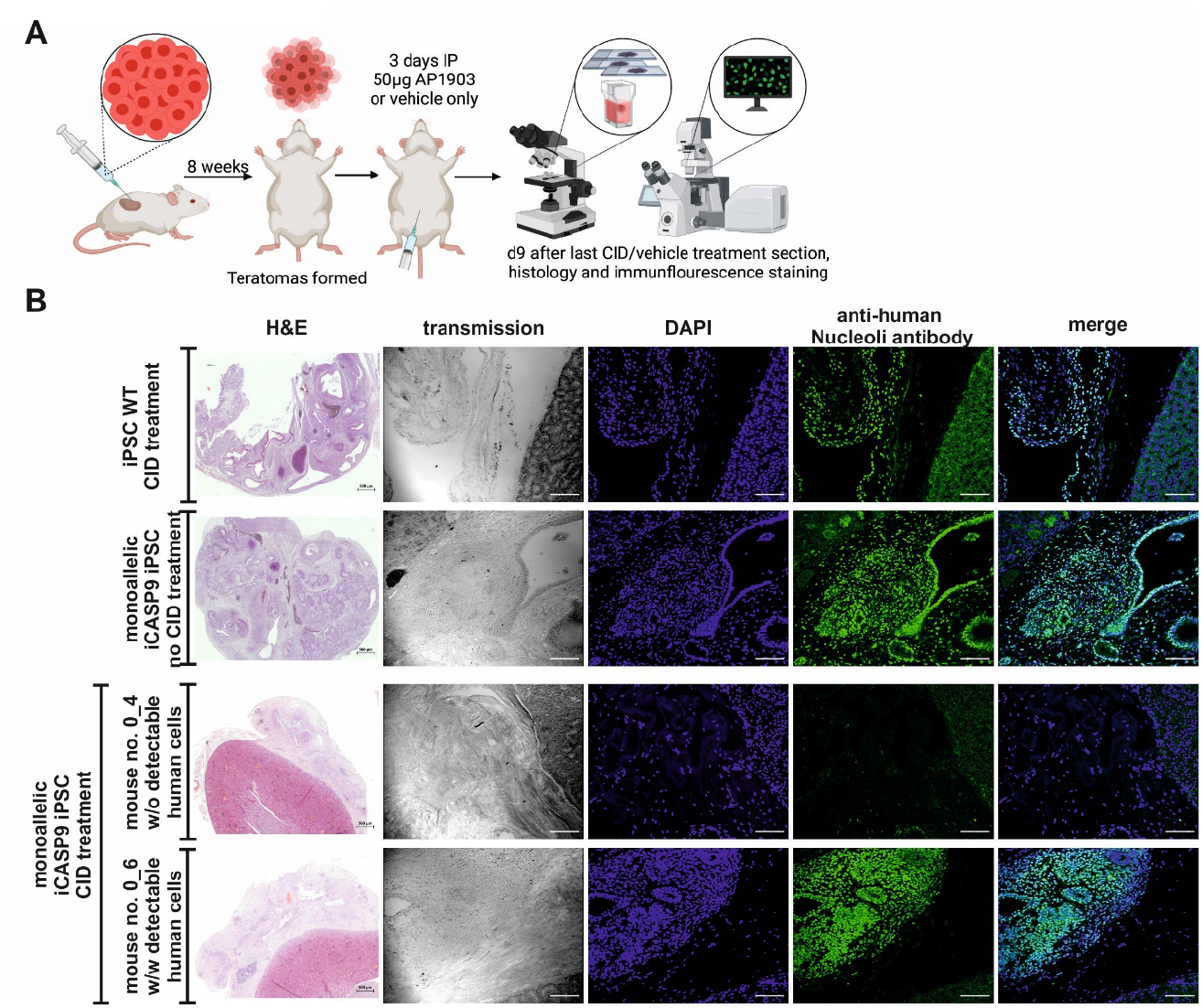
AP1903 eliminates preformed teratoma in the majority of NOD*SCID* mice after injection of monoallelic iCASP9 Phoenix iPSCs but did not prevent formation of human tumor-like tissue in all animals. A) Schematic illustration of the workflow for teratoma induction and CID treatment. B) NOD*SCID* mice were killed and sectioned 9 days after CID/vehicle treatment subsequent to injection of undifferentiated Phoenix iPSCs under the kidney capsule. (Immuno-)histology demonstrated growth of teratoma in all mice that received WT iPSCs and in monoallelic iCASP9 iPSCs without CID treatment (top and upper middle panels). In 3/5 mice that received iCASP9 iPSCs only some abnormal puffy mouse tissue with eosinophilic infiltration around the kidney could be detected (upper middle panel), probably representing fibrotic mouse tissue that formed in response to the massive CID-induced cell death and the resulting infiltration of phagocytes and granulocytes followed by pro-fibrotic cytokine release. Human tumor-like tissue was detected in 2/5 mice suggesting growth of CID-resistant cell clones (lower panel, see also table 1). Abbreviations: intraperitoneal, IP. (scale bars: 500μm for H&E, 100μm for other images)

**Table 1:** CID treatment did not reliably eliminate teratoma formed in NOD*SCID* mice after transplantation of monoallelic iCASP9 Phoenix iPSCs.

| iPSC clone | CID treatment<br>(50µg / d /<br>animal i.p.) | Number<br>of mice<br>treated* | Number of mice with detectable<br>human cell mass after CID-treatment |
| --- | --- | --- | --- |
| monoallelic iCASP9 | AP1903 | 5 | 2 |
| monoallelic iCASP9 | vehicle | 3 | 3 |
| wildtype | AP1903 | 5 | 5 |
| wildtype | vehicle | 3 | 3 |

\* 10<sup>6</sup> cells / mouse have been transplanted under the kidney capsule of NOD.CB17-PrkdcScid/J mice.

### *In vitro* CID treatment led to selection of rare CID-resistant cell subclones from monoallelic but not biallelic iCASP9 iPSC clones

In order to further explore the frequency of the observed CID resistance in monoallelic versus biallelic iPSCs, iCASP9 hCBiPS2 and iCASP9 Phoenix cells were treated with CID (concentrations from 0,5 nM-10nM AP1903 / AP20187), seeded onto irradiated feeder cells and were cultivated for three weeks to promote the propagation of potentially surviving cells. In correspondence to the *in vivo* experiments, survival of extremely rare CID resistant cell clones was observed in several independent experiments with overall frequencies of 3.6×10^-8^ for hCBiPSC or 2.5×10^-8^ for Phoenix iPSCs. We never observed surviving clones in biallelic iCASP9 iPSCs despite of the high total number of 0.8 billion treated cells (**Figure 4, table 2**).

**Figure 4:**
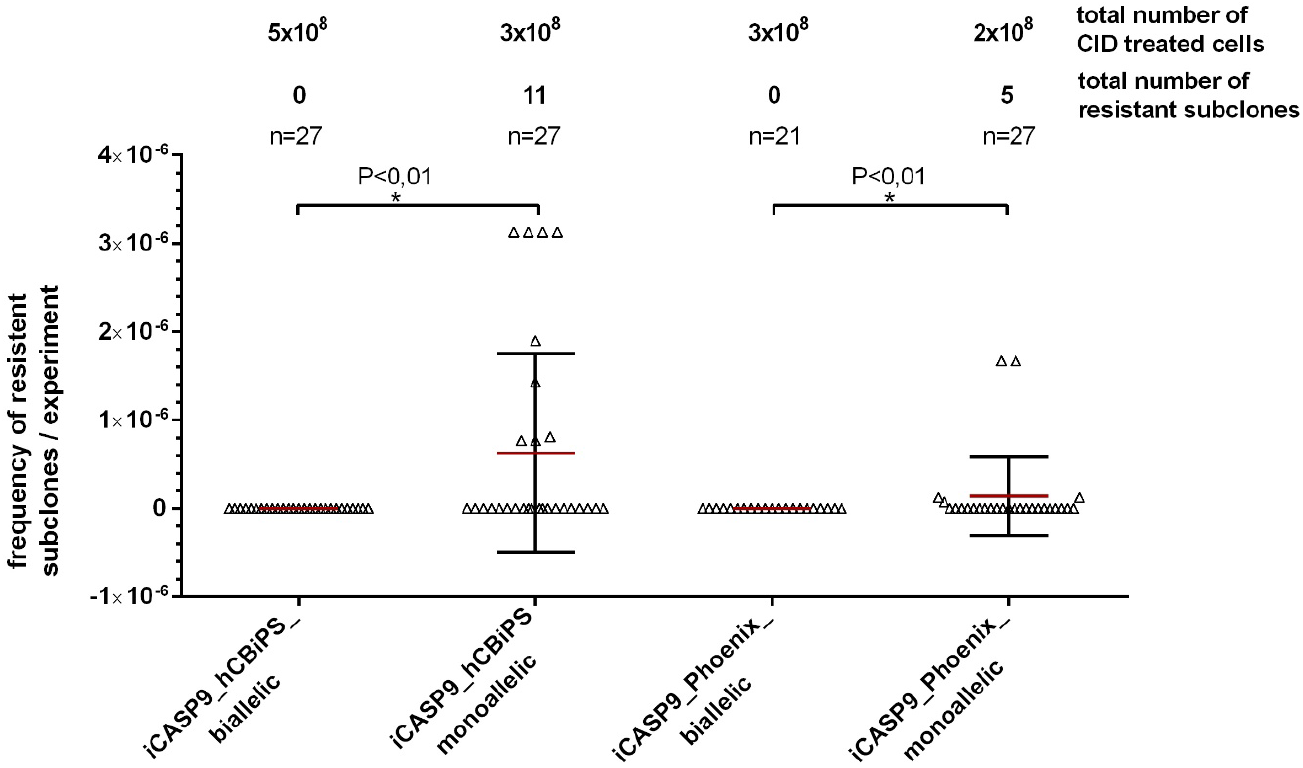
CID treatment of mono- and bialleic iPSCs led to selection of rare CID-resistant cell subclones from monoallelic but not biallelic iCASP9 iPSC clones. Monoallelic and biallelic iPSCs were treated with CID (concentrations from 0,5 nM-10nM AP1903 / AP20187), seeded onto irradiated feeder cells and were cultivated for three weeks to promote the propagation of potentially surviving cells. Survival of apparently CID resistant extremely rare cell clones was observed in several independent experiments (1×10^5^ to 1×10^8^ cells / experiment) with overall frequencies of 3.6×10^-8^ (hCBiPSC) or 2.5×10^-8^ (Phoenix). We never observed surviving clones in biallelic transgenic iPSCs despite of the high numbers of cell treated. Data are presented as mean ± SD (n = 21-27). D’Agostino & Pearson omnibus normality test. Kruskal-Wallis test (p<0,01).

**Table 2:** Rare monoallelic iCASP9 iPSC subclones become CID-resistant *in vitro* via loss-of-heterozygocity or methylation of the CAG promoter.

| iPS line | Number of CID-treated cells / experiment | Total number of CID-treated cells | CID conc. | Total number of CID-resistant subclones | Frequency of CID-resistant cell subclones | Subclones further cultivated and analysed | Subclones with eliminated transgene (confirmed/tested) | Subclones with CAG promoter methylated (confirmed/tested) |
| --- | --- | --- | --- | --- | --- | --- | --- | --- |
| Monoallelic iCASP9 hCBiPS2 | $1 \times 10^5$-$1 \times 10^8$ | $3 \times 10^8$ | 0,5nM-10nM | 11 | $3,6 \times 10^{-8}$ | 8 | 0/8 | 2/2* |
| Monoallelic iCASP9 Phoenix | $2 \times 10^5$-$8 \times 10^7$ | $2 \times 10^8$ | 1nM-10nM | 5 | $2,5 \times 10^{-8}$ | 4 | 3/4 | 0/1# |
| Biallelic iCASP9 hCBiPS2 | $2 \times 10^5$-$5 \times 10^8$ | $5,2 \times 10^8$ | 0,5nM-10nM | 0 | 0 | 0 | 0 | 0 |
| Biallelic iCASP9 Phoenix | $1 \times 10^5$-$3 \times 10^8$ | $3,14 \times 10^8$ | 1nM-10nM | 0 | 0 | 0 | 0 | 0 |

\* No further clones tested; # mechanism for CID resistance remained unclear

### Rare monoallelic iPSC subclones become CID-resistant due to transgene elimination, probably via Loss of Heterozygosity (LoH), or silencing via CAG promoter methylation

Twelve CID resistant subclones derived from monoallelic iCASP9 iPSCs (**Table 2**) were further cultivated to investigate mechanisms of the acquired CID resistance. Three out of four subclones derived from iCASP9 Phoenix did not show any dTomato expression. PCR based analysis of the genomic sequence of the transgene cassette in the AAVS1 locus of these iCASP9 Phoenix subclones (#1, #2 and #4 in **Figure 5B**) revealed elimination of the complete transgene, very likely via LoH. In the remaining clone with relatively weak dTomato expression neither transgene elimination nor increased methylation of CpGs in the CAG promoter or the adjacent AAVS1 (PPP1R12C) locus (**Fig S4**) as analyzed by Nanopore sequencing was observed (data not shown). In this case, the underlying mechanism remained elusive.

**Figure 5:**
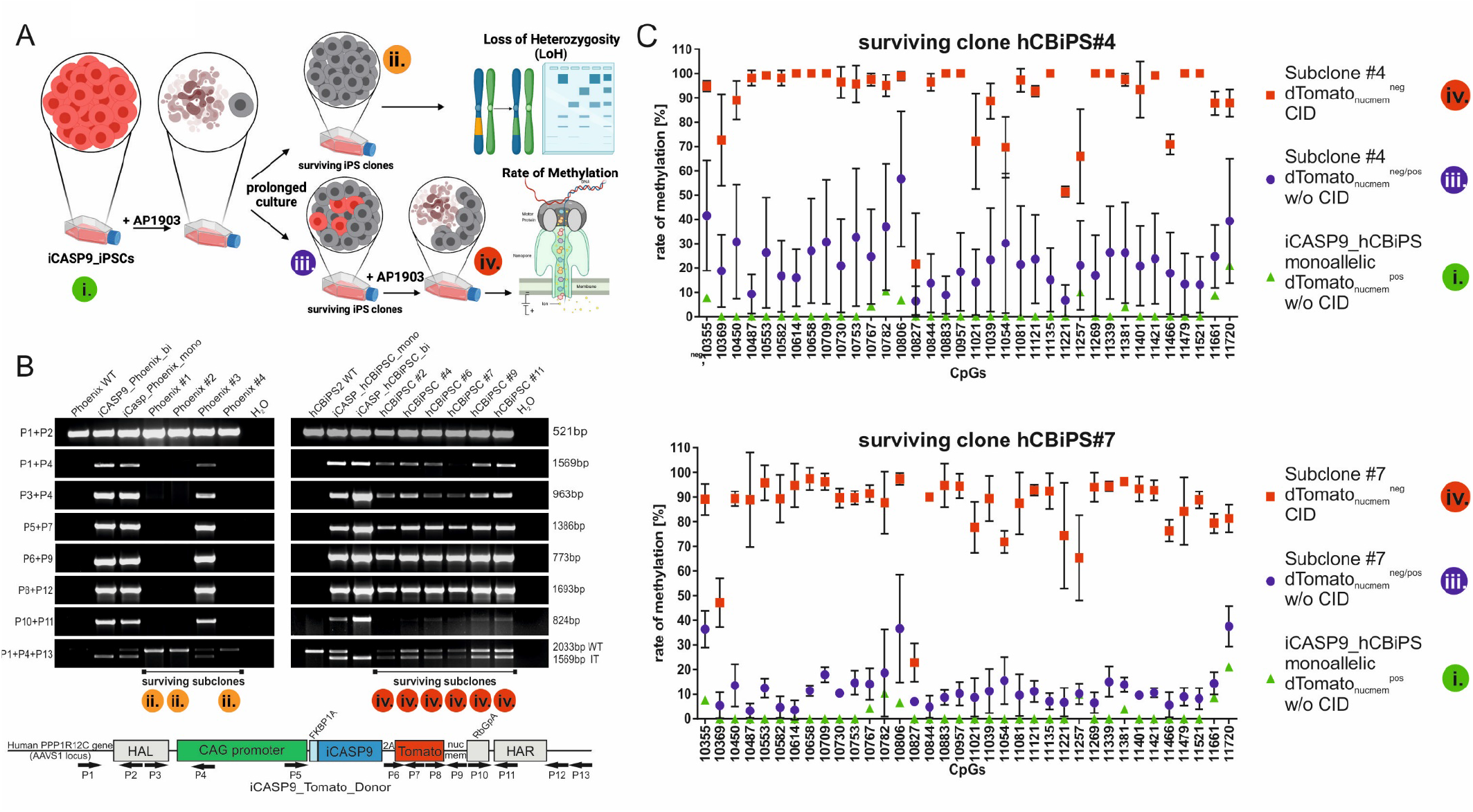
Rare monoallelic iPSC subclones become CID-resistant due to transgene elimination probably via Loss of Heterozygosity (LoH) or silencing via CAG promoter methylation. A) Scheme illustrating appearance and selection for Tomato^neg^ cells resistant to CID-induced apoptosis and re-appearance of Tomato^pos^ CID-sensitive cells during culture expansion, and analysis of different stages for LoH and methylation of promoter elements (and surrounding genomic DNA, data not shown). Colored circles mark stages that have been further analysed for LoH and promoter methylation. B) LoH occurred in three CID-resistant monoallelic iCASP9 Phoenix iPSC subclones but not in analyzed CID-resistant monoallelic iCASP9 hCBiPSC subclones (data not shown for hCBiPSCs). PCR based analysis of the genomic sequence of the transgene cassette in the AAVS1 locus revealed elimination of the transgene in Phoenix subclones #1, #2 and #4, very likely via LoH. Primer combinations and locations are depicted in the scheme below. C) Nanopore Sequencing showed methylation of the CAG promoter in 2 / 2 analyzed CID-resistant monoallelic dTomato_nucmem_^neg^ hcBiPSC subclones that did not show transgene elimination (data not shown). Analyses of CpG islands in the CAG promoter of surviving subclones #4 and #7 indicate strong correlation between cell survival and loss of nuclear dTomato expression after CID treatment, and high methylation rate in the CAG promoter. Data are presented as mean ± SD (n = 3-4). A scheme of CpG islands in the AAVS1 locus and the integrated iCASP9 donor construct is shown in Figure S4. No elevated methylation of the adjacent AAVS1 (PPP1R12C) locus was observed (data not shown).

Loss of the transgene could not be demonstrated in any of the eight analyzed iCASP9 hCBiPS2 subclones. Interestingly, repeated CID-treatment led to elimination of dTomato expressing cells in all these CID-resistant subclones, however, dTomato expressing cells re-appeared in all subclones several days after CID expression (**Figure S3**) suggesting reversible epigenetic mechanisms to be responsible for transgene silencing.

Exemplarily, two of these phenotypically similar subclones (#4 and #7) were analyzed for increased methylation of CpGs in the AAVS1 locus or the CAG promoter via Nanopore sequencing. The observed high methylation rate of the CAG promoter in these subclones (**Figure 5C**) perfectly correlates with loss of dTomato expression and their resistance towards CID-treatment. Again, no elevated methylation of the adjacent AAVS1 (PPP1R12C) locus was observed (data not shown).

## Discussion

The availability of human ESCs and iPSCs with their far-reaching potential for proliferation and differentiation offers novel opportunities for the development of tailored cellular therapies. Therapeutic risks associated with PSC-based therapies include ones that are related to abnormal cell functions such as arrhythmia in case of cardiomyocytes or dysregulation of hormones in pancreatic islands or dopaminergic neurons, but especially tumor formation is a major concern. Integration of synthetic fail-safe systems into PSCs have been proposed as additional safety measure enabling induced suicide in case of transplant-associated tumor formation. Whereas in case of retro- and lentiviral delivery of such systems, the appearance of suicide-resistant subclones is not surprising considering the frequently observed high levels of transgene silencing, targeted integration into genomic loci known for their robust expression was considered as more appropriate approach to achieve reliable induced elimination of transplanted cells. A recent study, however, challenged that estimation. With a total of eight suicide-resistant escapee clones among a total of 120 million cells for a monoallelic HSV-TK suicide gene integrated into the endogenous CDK1 locus, Liang et al. observed an overall frequency of 6.7×10^-8^ escapees ^6^.

With a total of 16 escapee clones among 500 million cells (3.2×10^-8^) we have now observed a comparable frequency for the CAG-iCASP9 safety switch inserted in the AAVS1 locus. Similar to Liang et al., we never observed resistant cell clones that develop from iPSCs carrying a homozygous (biallelic) safety switch. This finding further argues for LoH as underlying mechanism of transgene elimination in monoallelic subclones, since elimination of the suicide gene via LoH can be excluded in cells homozygous for the integrated safety switch.

In case of the observed aberrant promoter methylation as underlying mechanism for safety switch inactivation the question remains whether such rare events depend on the status of the cellular methylation machinery, which would imply that in the event of methylation of one allele there is also an increased likelihood for similar methylation of the second allele. The fact, however, that despite a high number of 0.8 billion cultured iPS cells we never observed any escapee from two iPSC lines with integrated biallelic iCASP9 safety switch suggest that the observed aberrant methylation of the promoter that occurs in very rare cases (~3×10^-8^) on one allele is completely independent of the second allele. If this hypothesis applies, a simultaneous methylation of both alleles would be extremely unlikely (3×10^-8^ × 3×10^-8^ = 9×10^-16^).

In theory, also individual loss of function mutations may lead to CID resistance. Homologous recombination may even lead to transmission of the mutation to the 2^nd^ allele. While such events are also considered very rare and were not observed in our experiments, they certainly deserve further research. Also further mechanisms such as gene silencing due to histone modifications have to be considered.

Our results indicate that even targeted integration of a monoallelic safety switch into a safe harbor site known for robust transgene expression is insufficient to exclude development of CID-resistant subclones. Although a biallelic approach most likely offers much higher safety levels, the question remains how to calculate the safety level not only for a specific cell product but also for a given clinical scenario. Addressing that question, Liang et al. developed the term *safe-cell level (SCL)* as *the number of therapeutic batches in which there is expected to be one none-safe batch*^6^. This SCL, which is actually reflecting the frequency of escapees, was then put in relation to clinically relevant cell numbers in terms of the therapeutically applied cell doses. These will often be substantially higher than typical cell numbers handled in conventional cell culture systems and may range from 10^5^ to 10^10^ cells, e.g. it is estimated that ~ 10^9^ cardiomyocytes would have to be replaced after myocardial infarction.

It is debatable, however, whether relation of a calculated SCL to the size of a required therapeutic cell dose is the most appropriate approach to estimate the safety level of a given cell product. Integrated suicide genes are mainly considered as safety measure to eliminate proliferating tumor cells that either arise as teratoma from contaminating undifferentiated pluripotent stem cells, or develop from rare cell clones that carry mutations, which lead to altered expression or function of oncogenes. It is, however, extremely unlikely that a very rare cell clone within the therapeutic cell batch that acquired CID resistance either through LoH, silencing of the transgene or other randomly developed mutation, undergoes another very rare event, which is tumor transformation.

More relevant are other scenarios, where *in vivo* single contaminating undifferentiated PSCs among the therapeutic cell transplant form a teratoma, or an individual cell clone acquires (a) genetic aberration(s) and undergoes tumor transformation, in both cases followed by massive proliferation. Among the generated large number of tumor cells, clonal CID resistance may develop as a second, independent and again very rare, event.

Therefore, instead of calculating a safe-cell-level in consideration of the size of a produced cell batch or the therapeutic cell dose, it seems more reasonable to estimate the safety of a cell product by putting the frequency of suicide-resistant escapee clones in relation to the number of cells in a tumor mass detectable during routine tumor screening.

Actually, recent estimations for cell numbers in tumors tissue range from 10^7^-10^8^ cells for a tumor of 1 cm^3^, a size that should be reliably detectable by modern imaging approaches ^14^. Large tumors may even contain more than 10^9^ to 10^10^ cells. Provided a frequency of ~ 5×10^-8^ for clonal suicide escapees in cells with a monoallelic safety switch and 10^8^ cells in a tumor, it is obvious that it is not unlikely that a tumor when it becomes large enough to become detected clinically may already contain CID-resistant tumor cell clones.

In light of these considerations, monoallelic integration of synthetic safety switches clearly appears insufficient to provide an increased safety level for cell therapy products. In contrast, we could not detect any drug resistant escapees from biallelic iCASP9 hiPSCs among 0.8 billion iPSCs. That cell number is already ~10x higher than the estimated cell number in a tumor of 1cm^3^ in size, which should be reliably detectable by regular tumor screening e.g. via MRI, before further increased tumor cell numbers may lead to escapees. Large mass production settings would be required to further define the actual risk for appearance of clonal escapees from cells with integrated biallelic safety switches. Experimental proof, however, of the hypothesized independency of suicide transgene inactivation on both alleles, which would imply extremely low frequencies of clonal escapees in a range of 10^-15^ is almost impossible. Since even with most advanced mass culture technologies production of not more than 10^7^ cells/ml is possible ^15^, a culture volume of approximately 100.000l would be required to generate the huge number of 10^15^ cells.

## Material and Methods

### Plasmid construction

The used AAVS1 TALENs were generated via the golden gate assembly method {Morbitzer, 2011 #48} and contain wild-type FokI nuclease domains. The AAVS1 locus-specific TALEN expression cassettes ^16^ were placed under control of a CAG promoter.

The AAVS1.iCaspase9 donor was based on the vector SFG.iCasp9.2A.ΔCD19 described in Stasi et al 2011. For generation of the AAVS1 donor plasmid standard cloning technologies were used. The AAVS1 targeting vector contains CAG-iCasp9-2A-dTomato_nucmem_-RbGpA flanked by two arms of ~700 bp AAVS1 locus homology sequences in a pUC19 expression vector backbone (Thermo Fisher Scientific).

As described in ^5^, the transgene iCasp9 consists of the sequence of the human FK506-binding protein (FKBP12; GenBank number, AH002818) with an F36V mutation, connected through a Ser-Gly-Gly-Gly-Ser linker to the gene encoding human caspase 9 (CASP9; GenBank number, NM001229).

### Cell culture

We used two human iPSC (hiPSC) lines generated in our group. hCBiPS2 is based on the lentiviral transduction of cord-blood-derived endothelial cells {Haase, 2009 #50} and Phoenix is based on cord-blood-derived CD34+ cells transduced with sendai virus vectors {Haase, 2017 #56}. The hiPSC lines were cultured and expanded on irradiated mouse embryonic fibroblasts (MEFs) in iPS medium (knockout Dulbecco’s modified Eagle’s medium supplemented with 20% knockout serum replacement (KSR), 1 mM L-glutamine, 0.1 mM b-mercaptoethanol, 1% nonessential amino acid stock (all from Thermo Fisher Scientific), and 10 ng/ml basic fibroblast growth factor (bFGF; supplied by the Institute for Technical Chemistry, Leibniz University, Hannover, Germany) {Chen, 2012 #51}).

### Transfection and clone establishment

For Nucleofection hiPSCs were expanded as monolayer cultures on Geltrex (Thermo Fisher Scientific), cultivated in mTESR1 (STEMCELL Technologies) and harvested by Accutase (Thermo Fisher Scientific). The transfection was performed with Neon^®^ Transfection system (ThermoFisher Scientific). 1×10^6^ cells were resuspended in 105 μL Neon^®^ buffer, electroporated with 5 μg of each plasmid encoding for AAVS1-specific-TALEN and 15 μg donor plasmid with two pulses at 1,000 V for 20 ms, and plated onto Geltrex-coated dishes with MEF conditioned medium (CM) (DMEM/F12 supplemented with 15% KSR, 100μM β-mercaptoethanol, 1% nonessential amino acid stock (all from Thermo Fisher Scientific), 10ng/mL bFGF and 10 mM Y-27632 (both from the Institute for Technical Chemistry, Leibniz University Hannover).

Transfected cells were cultivated as a monolayer prior to Fluorescence Activated Cell Sorting (FACS). For clone generation, cells were harvested from the monolayer culture by Accutase on day 10 after transfection, and were sorted on the FACSAria IIu (BD Bioscience) or XDP (Beckman-Coulter) for of dTomato_nucmem_ positive cells. Sorted populations were plated at low density onto Geltrex-coated dishes. Colonies were picked manually and transferred into feeder-based culture conditions resulted in single-cell clones, which were also analyzed via PCR screening (see schematic illustration in Figure S1).

### PCR Screening and characterization of transgenic clones

Genomic DNA was prepared using the QIAamp DNA Blood Mini Kit (QIAGEN) according to the manufacturer’s instructions, and 100ng of gDNA was amplified by PCR with GoTaq DNA polymerase (Promega). Sequences and specifications of the primers are shown in Table S1.

### Karyotype analysis

After treatment of adherent hiPSC for 30 min with colcemid (Invitrogen), cells were trypsinised and metaphases were prepared according to standard procedures. Fluorescence R-banding using chromomycin A3 and methyl green was performed as previously described in detail (Schlegelberger et al., 1999). At least 15 metaphases were analyzed per clone. Karyotypes were described according to the International System for Human Cytogenetic Nomenclature (ISCN).

### Immunohistochemistry for characterisation of transgenic clones

Cells were fixed with 4% paraformaldehyde (w/v), permeabilized with triton blocking solution and stained by standard protocols using primary antibodies, listed in Table S2, and appropriate secondary antibodies. Incubation with primary antibodies and corresponding isotype controls was performed overnight at 4 °C. Staining of living cells for extracellular markers was performed for 1 h. Secondary antibody staining was performed afterwards. Cells were counterstained with DAPI (Sigma) for the analyzes with an AxioObserver A1 fluorescence microscope and Axiovision software 4.71 (Zeiss) or Propidium Iodid (PI) (final concentration 1μg/ml) (Thermo Fisher Scientific) for the analyses using a MACSQuant^®^ Analyzer (Miltenyi Biotech). Flow cytometric data evaluation was done with FlowJo 7.6.5 software (Celeza).

### *In vitro* differentiation

HiPSC cultured on MEFs in iPS medium were detached with CollagenaseIV (Thermo Fisher Scientific), dispersed in small clumps and cultured in differentiation medium (80% IMDM supplemented with 20% fetal calf serum, 1mM L-glutamine, 0,1mM ß-mercapthoethanol and 1% nonessential amino acid stock) in ultra-low-attachment plates (Corning Inc., NY, USA) for 7 days. On day 7 the formed embryoid bodys (EB) were plated onto 0,1% gelatin coated tissue culture dishes and cultured for a further 13 days before fixation and immunostaining.

### Induction of apoptosis with the Chemical Inducer of Dimerization AP20187 and AP1903

iPSCs were seeded on Geltrex-coated tissue culture plates, cultivated in mTESR1 and grown 48hrs before functionality of the suicide gene was assessed by adding the chemical inducer of dimerization (CID) (AP20187, B/B Homodimerizer, Clontech Laboratories, Inc), or AP1903. AP1903 was synthesized in house at the Institute for Technical Chemistry, Leibniz University, Hannover, Germany. CIDs were applied for 24 h with concentrations of 0,001nM, 0,1nM, 1nM, and 10nM. For flow cytometric analysis, iCASP9_hiPSCs not treated with CID or the hiPSC wildtype were dissociated into single cells using Accutase. Samples based on iCASP9_hiPSC treated with CID in different concentrations were already in suspension.

For detection of living cells undifferentiated hiPSCs were stained with 1μmol Calcein AM (Thermo Fisher Scientific) for 45 minutes and additionally all samples were in parallel analyzed using PI. Data were acquired on MACSQuant^®^ Analyzer and analyzed using FlowJo software (Flowjo, LLC).

### Teratoma assay

For in vivo testing of iCASP9_suicide switch functionality, the iCASP9_hiPSCs were cultivated as colonies on MEFs in iPS medium. After detachment with CollagenaseIV the cell suspension was mixed in ratio 1:1 with matrigel (Thermo Fisher Scientific) to increase cell survival. Teratomas were induced by injection of the cell/matrigel suspension (~1 million cells per injection) under the renal capsule of NOD/*SCID* mice. After 8–10 weeks, the animals showed increased abdominal girth and body weight.

Then, the mice were administrated 50μg/day/animal (2,5mg/kg) AP1903 (4% AP1903 stock solution (3,125mg/ml), 10% PEG 400 (100%), and 86% Tween (2% in water)) or the same amount of PEG400/Tween solution only as vehicle. The injection was given intraperitoneal (IP) for three days. Nine days after last injection mice were sacrificed. During all experiments, the ‘‘Principles of laboratory animal care’’ (NIH publication No. 86-23, revised 1985) as well as the Animal Welfare Law of Lower Saxony were followed.

### Histological Analysis and Confocal microscopy

For histological analyses, the samples were fixed in formalin and processed for paraffin embedding according to standard protocols. Histological staining was conducted on paraffin cuts using hematoxylin and eosin (H&E).

For immunofluorescence analyses of human cells, tissue slides of paraffin were used. Antigen retrieval was achieved by preheated sodium citrate buffer according to standard protocols. After antigen retrieval, tissue slides were stained using the protocol described before for immunohistochemistry.

Confocal images were acquired with a Zeiss LSM 780 and ZEN black 2.3. To separate the immunostaining signal from autofluorescence background online fingerprinting mode was used. For that reference spectra were taken from single fluorescence staining. Several autofluorescence reference spectra were taken from specific regions of non-stained samples and used for unmixing. Images were obtained with a 20×/0.8 Plan-Apochromat, 405 nm and 488 nm excitation at zoom 0.6 (image dimensions 1024 × 1024 pixel, 700 × 700 μm).

To correctly compare immunostainings all samples were prepared in parallel and imaged at identical acquisition settings. Images were displayed at the same brightness settings to emphasize the specificity of the conditions. A faint gauss blur was applied to the images.

### *In vitro* testing for CID treatment surviving subclones

IPSCs were seeded on Geltrex-coated tissue culture plates or flasks, cultivated in mTESR1 and grown until a confluent monolayer is reached. Medium was changed to medium containing the chemical inducer AP20187 or AP1903 in concentrations from 0,5nM up to 10nM. After 24 hours, the complete medium containing all treated cells were centrifuged and transferred into feeder-based culture conditions. The treated hiPSCs were cultured in iPS medium (described in Cell Culture). In the first 24 hours the medium was supplemented with 10 mM Y-27632 to allow the survival of single cells. After 3 weeks of cultivation surviving colonies were picked, cultivated further under feeder-based culture conditions. After prolonged cultivation surviving subclones were transferred to feeder-free-conditions described before and were treated again with 10nM AP1903 (see also illustration Figure 5). The achieved population was analyzed for rate of methylation via Nanopore Sequencing and as described before (PCR Screening) for correct integration of the iCASP9-Donor construct (illustration Figure 5). Sequences and specifications of the primers are shown in Table S1.

### Nanopore sequencing

Genomic DNA samples were collected before and after the treatment with AP1903 and stored as a pellet at minus 80°C. Isolation of genomic DNA was done using the NucleoBond^®^ HMW DNA Kit (Macherey Nagel) according to the manufacturer’s instructions. For Cas-mediated PCR-free enrichment of the construct integrated between base position 55115763 and 55115769 on chromosome 19 of hCBIPSC and Phoenix IPSC guideRNAs were designed by means of the “Alt-R Custom Cas9 crRNA Design Tool” (IDT^®^). For this purpose, the sequences of the regions, in which the potential guide RNAs should be located in order to obtain fragments of approximately 5.000 base pairs length, were pasted into the dialogue box. The program outputs alternative guide RNAs in the inserted region and indicates the respective on-target potential (values from 0 to 100, the higher the better) and off-target risk (values from 0 to 100, the higher the lower the off-target risk). Only guides with an on-target-potential ≥ 60 and an off-target risk ≥ 75 were accepted for our sequencing experiments and were ordered from IDT (Integrated DNA Technologies, Coralville, USA). During establishment of the best guideRNA strategy, seven guide RNAs were tested. However, according to the criteria how guideRNAs should be positioned in relation to the target (for more detail see also https://biorxiv.org/cgi/content/short/2021.09.17.460763v1), a combination of the guideRNAs no. 2134, 2136, 55 and 2139 (tableS3) proved to be most effective as it yielded the highest number of calls per CpG-side. It is of utmost importance to obtain a sufficient number of calls per CpG-site (https://biorxiv.org/cgi/content/short/2021.09.17.460763v1) in order to be able to make a valid statement about the methylation rates of the CpGs. The respective sequences of the guides and their position in the construct are shown in figure S4. To enable sequencing of this CG-rich and therefore difficult to sequence area, the DNA had to be cut into shorter pieces of about 5.000 base pairs each in order to avoid the formation of secondary structures. Before sequencing, the DNA quality was assessed in order to ensure that only high-molecular weight DNA was used in the subsequent library preparation for the Nanopore run. For this purpose, a pulsed-field gel analysis was carried out using a Pippin Pulse electrophoresis power supply (Sage Science, Beverly, USA). 5 μg of DNA were used per sample for the library preparation by the Cas-mediated PCR-free enrichment protocol. The latter includes the usage of the Ligation Sequencing (SQK-LSK109) Kit (Oxford Nanopore Technologies, Oxford, UK) and the Native Barcoding Expansion 1-12 (PCR-free) (EXP-NBD104) Kit (Oxford Na-nopore Technologies, Oxford, UK). The entire process consists of the following steps: Dephosphorylation of genomic DNA, Preparation of the Cas9 ribonucleoprotein complexes (RNPs), Cleavage and dA-tailing of target DNA, Native barcode ligation with appropriate purification by beads and final pooling of samples, Adapter ligation (due to barcoding, AMII has been used instead of AMX) and purification by AMPure XP beads (0.3 × volume). The pellet was resuspended in 14 μl of preheated elution buffer at 37°C for 20 minutes and 12 μl of the eluat were added to sequencing buffer and loading beads for the final library preparation step. Afterwards, the SpotON flow cell FLO-MIN106D (Oxford Nanopore Technologies, Oxford, UK) was primed and loaded, and the sequencing run was started on a MinION-device (Oxford Na-nopore Technologies, Oxford, UK).

For basecalling and demultiplexing data guppy basecaller v5.0.11 (Oxford Nanopore Technologies, Oford, UK) was used with standard settings and config file ‘dna_r9.4.1_450bps_hac’. The basecalled reads were aligned to the reference genome using minimap2 v2.20 (https://doi.org/10.1093/bioinformatics/bty191). For methylation calling we used nanopolish v0.13.2 (DOI: 10.1038/nmeth.4184).

### PCR-based analysis of Loss of Heterozygosity (LoH)

Genomic DNA was prepared using the QIAamp DNA Blood Mini Kit (QIAGEN) according to the manufacturer’s instructions, and 100ng of gDNA was amplified by PCR with GoTaq DNA polymerase (Promega). Sequences and specifications of the primers are shown in Table S1.

### Statistical analyses

Data are presented as mean ± SD (n = 21-27) and were tested for normal distribution using theD’Agostino & Pearson omnibus normality test. As data points were not normally distributed, group comparison was performed by means of the non-parametric Kruskal-Wallis test (p<0,01) for independent samples.

## Acknowledgements

The authors thank Jennifer Beier, Janina Zöllner, Nicole Cleve, Annika Franke, Alexandra Lipus and Viktor Lutscher for technical assistance and helpful discussion. Illustrations in figure 1, 3 and 5 were created with BioRender.com.The plasmid SFG.iCasp9.2A.ΔCD19 was kindly provided by G. Dotti, Baylor College of Medicine. This work was supported by the Cluster of Excellence, REBIRTH (EXC 62/1, 62/3).

## Author Contributions

S.W. designed, performed and analysed experiments. S.M. and A.H. was involved in the cloning procedure of the donor. and design of guide RNAs. K.J. and M.D. designed, performed and analysed Nanopore Sequencing. S.G. analysed the H&E stainings and provided expertise and feedback. W.K. and A.M. performed and analysed the *in vivo* experiments (teratoma assay). A.K. provided Y-27632 and AP1903. A.Z. and E.P. were involved in analyses of remaining human cells and edit the resulting pictures (confocal microscopy). G.G. performed and analysed the karyotype analysis. M.A. provided expertise and feedback. N.L. helped with data interpretation. T.M. contributed to experimental design. R.Z. contributed expertise and helpul discussion. U. M. and S.W. wrote the manuscript.

## Supplement

### Supplementary Figures

**Figure S1:**
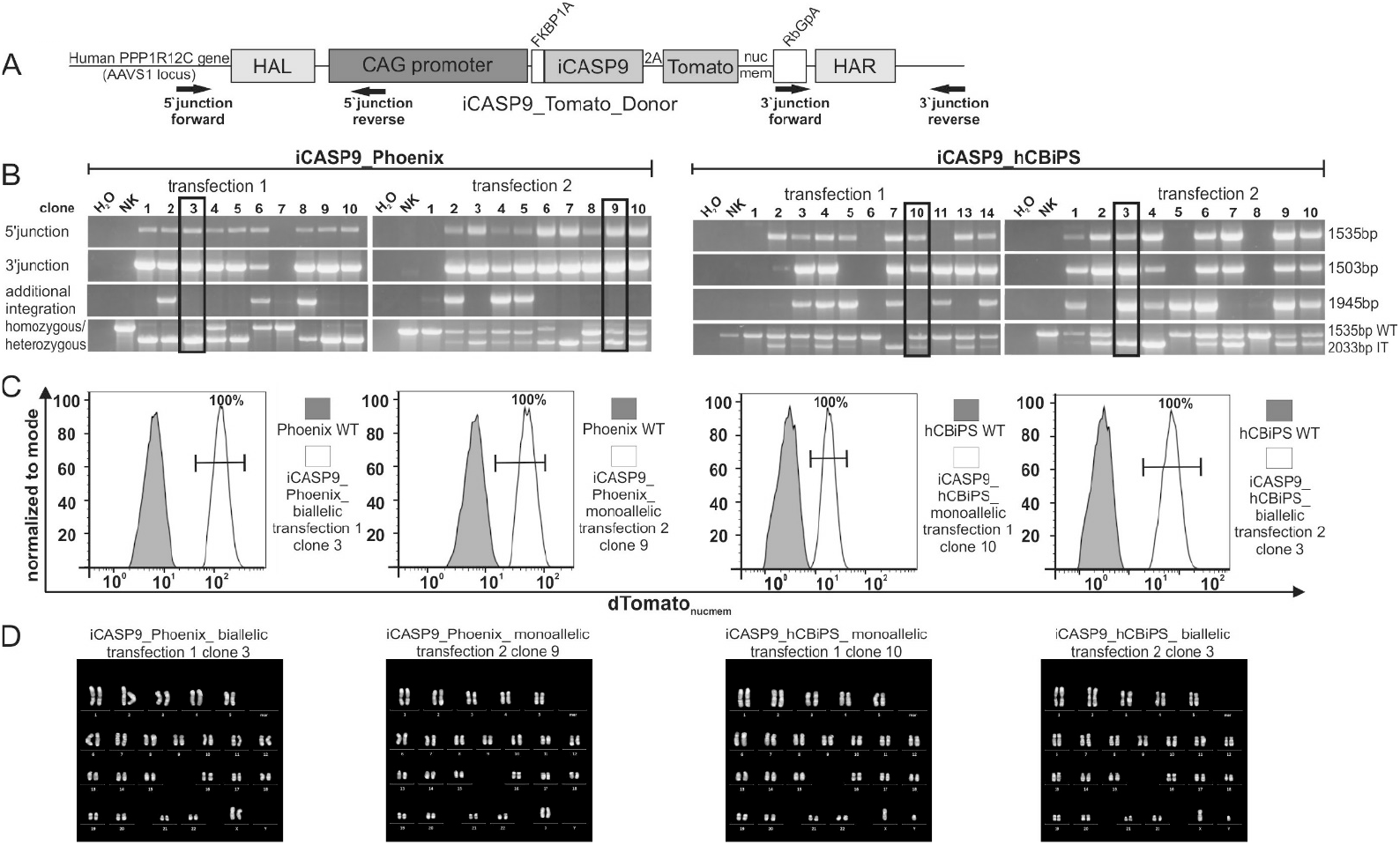
Four iCASP9 iPSC clones were selected for further studies based on genotyping and flow cytometric analysis of dTomato_nucmem_ expression and karyotype analysis. A) Schematic illustration of Primer combinations and locations used for genotyping of targeted iPSCs. B) Correctly targeted and dTomato_nucmem_ ^pos^ clones (marked with frames) were preselected based on demonstration of mono-/biallelic integration by junction PCRs on genomic DNA using primer pairs spanning the 5’- and the 3’-junction of the donor cassette and genomic AAVS1 sequence. Expression of dTomato_nucmem_ was analyzed via flow cytometry. Histograms for dTomato_nucmem_ expression of transgenic clones are shown exemplarily. Karyotype analysis demonstrated absence of larger genomic aberrations. Representative metaphases are shown. Chromosome analysis was performed following standard cytogenetic procedures. At least 20 metaphases per clone were analysed.

**Figure S2:**
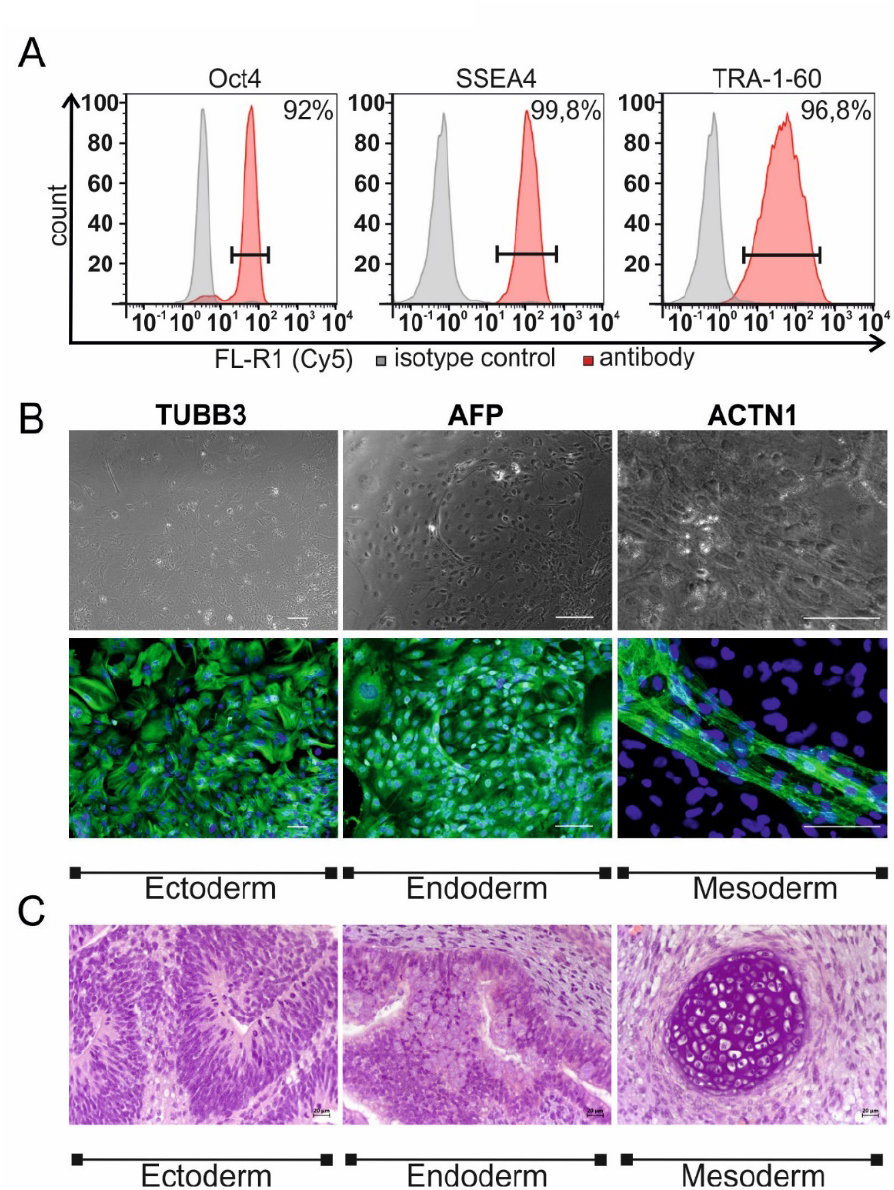
Characterization of iCASP9_hiPSCs (exemplarily shown for iCASP_Phoenix_monoallelic). A) Immunofluorescence staining of iCASP9_hiPSCs against OCT4, SSEA-4 and TRA-1-60 and analyzes via flow cytometry demonstrate the expression of these typical pluripotency markers. B) Immunostaining of iCASP9_hiPSCs derivatives on d21 of differentiation revealed expression of ectodermal (TUBB3), endodermal (AFP), and mesodermal (ACTN1) marker proteins (green). Nuclei are stained with DAPI (blue). Scale bars: 100 μm. C) Injection of undifferentiated iCASP9_Phoenix_monoallelic iPSCs into immunodeficient NOD*SCID* mice led to formation of teratomas containing derivatives of all three germ layers. Neural tube formation representing ectodermal differentiation. Endodermal epithelium with prominent mucus-producing cells representing endoderm and mesoderm formation. Chondrocytes showing mesoderm formation. (Scale bars represent 20 mm as depicted.)

**Fig. S3:**
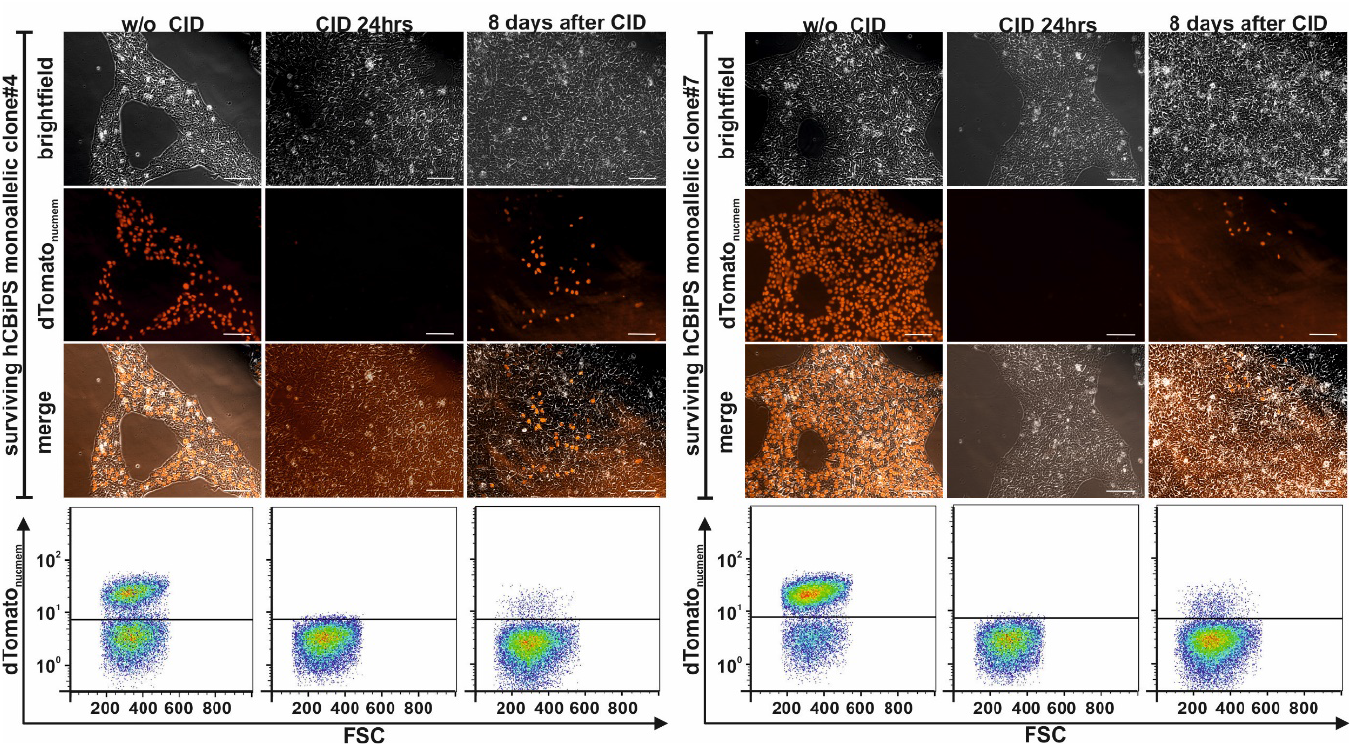
dTomato_nucmem_^neg^ colonies that survived CID treatment partially regain dTomato_nucmem_ expression after prolonged cultivation. Representative microscopic pictures and flow cytometrical analyses of two monoallelic iCASP9 hcBiPS2 subclones treated with CID are shown. Left panel: monoallelic iCASP9 subclone #4, right panel: monoallelic iCASP9 subclone #7. Left columns: show monoallelic hcBiPS single cell clones that survived initial treatment with CID during recovery under feeder-based culture conditions. These cultures contain dTomato_nucmem_^pos^ and dTomato_nucmem_^neg^ cells (Stage iii in Figure 5A). Middle colums show the hcBiPSC clones, treated for 24 hours with CID to eliminate the dTomato_nucmem_^pos^ fraction. Surviving cells are dTomato_nucmem_^neg^ (Stage iv in Figure 5A). Starting eight days after CID-treatment new dTomato_nucmem_^pos^ cells could be observed (scale bar 100μm).

**Fig. S4:**
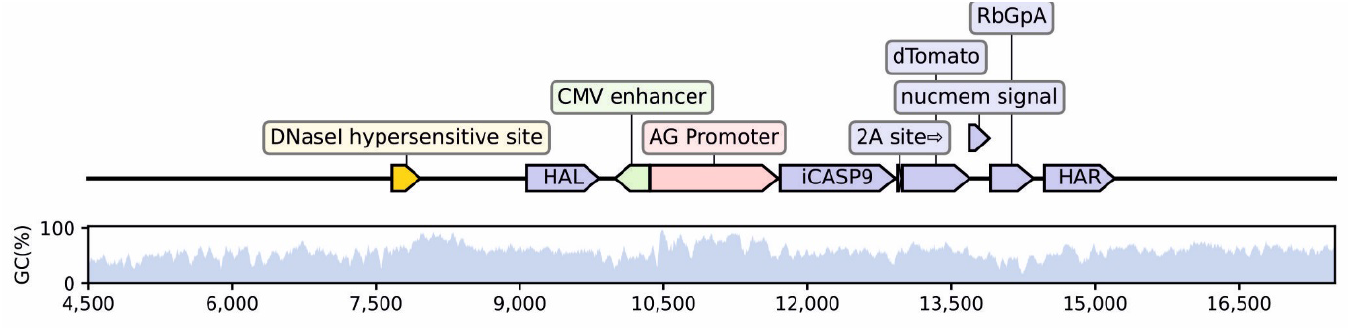
Schematic illustration of CpG contents in the iCASP9_Donor integrated into the AAVS1 locus. Content of CpGs is shown for the iCASP9_Donor and surrounding areas of the human PPP1R12C gene.

### Supplementary Tables

**Table S1:**
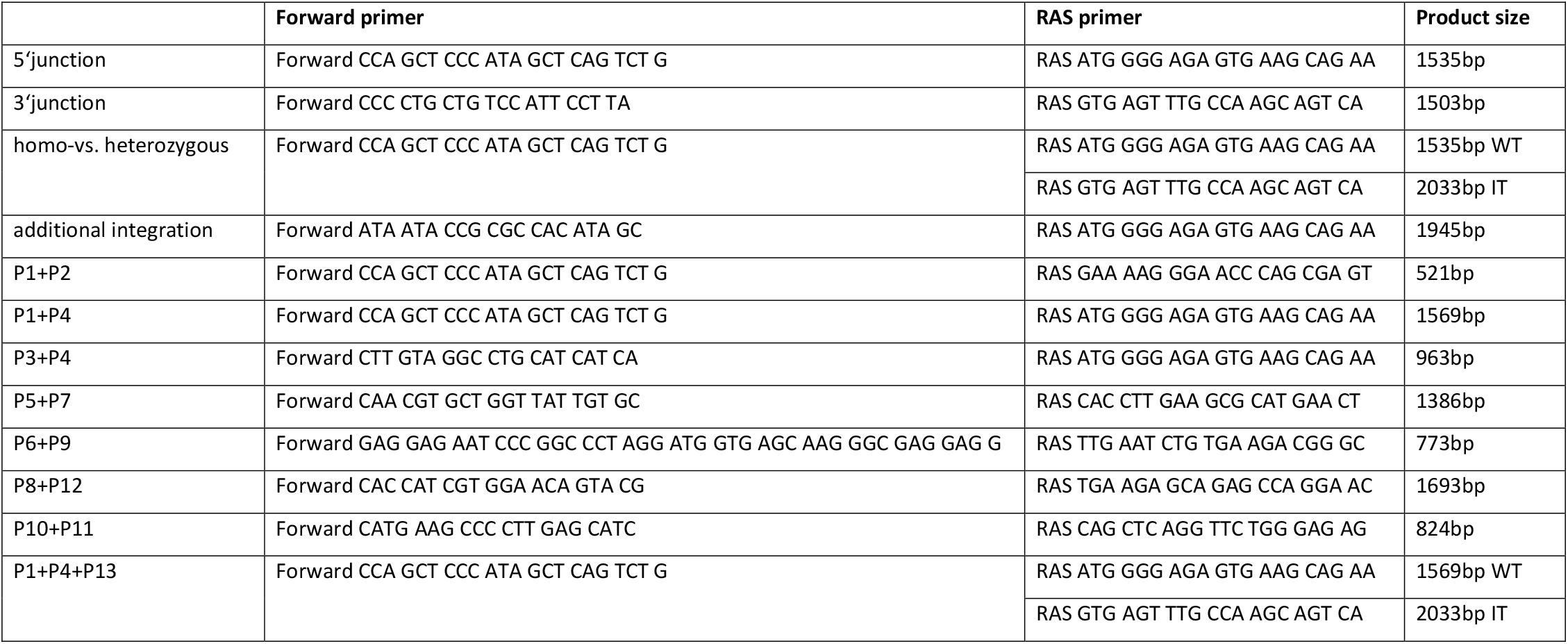
Primers and expected sizes of PCR products. Abbreviations: IT, Integrated transgene; RAS: Reverse antisense; WT, Wild type.

**Table S2:** Antibodies used for immunocytochemistry / flow-cytometry.

|  | **Antibody** | **Dilution** | **Company Cat # and RRID** |
| --- | --- | --- | --- |
| **Pluripotency Markers** | Mouse anti-OCT4 | 1:100 | Santa Cruz Cat# sc-5279 |
|  | Mouse anti-TRA-1-60 | 1:100 | Abcam Cat# 16288 |
|  | Mouse anti-SSEA-4 | 1:100 | DSHB Cat# MC813-70 (ab16287) |
| **Differentiation Markers** | Mouse anti-α-Fetoprotein | 1:300 | R&D Cat# MAB1368 |
| | Mouse anti-α-Actinin, Sarcomeric | 1:800 | Sigma Aldrich Cat# A7811 |
| | Mouse anti-β-3-Tubulin | 1:400 | Millipore Cat #05-559 |
| Detection of human cells | Mouse anti-human-Nucleoli | 1:400 | Abcam Cat# ab190710 |
| Secondary antibodies | Cy5 Donkey anti-mouse IgG | 1:200 | Jackson ImmunoResearch Labs Cat# 715-175-150, |
|  | AF488 Donkey anti-goat IgG | 1:200 | Jackson ImmunoResearch Labs Cat# 705-545-147 |

**Table S3:**
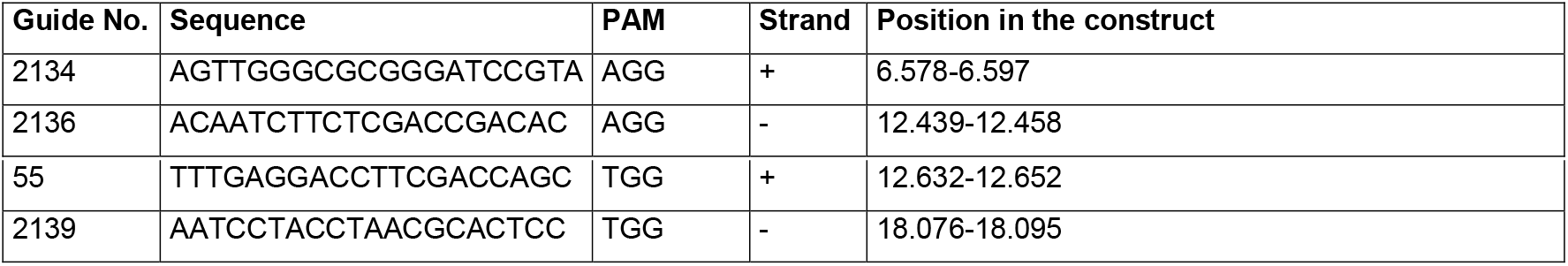
Overview of the GuideRNAs used for the Cas-mediated PCR-free enrichment of the construct sequence for Nanopore sequencing, including the respective sequences, PAMs (Protospacer Adjacent Motifs), strand information and position in the construct.

